# Modulation of adaptive immune responses by *Akkermansia muciniphila* is restricted to an early life window in NOD mice

**DOI:** 10.1101/2022.08.16.504185

**Authors:** Sarah Maddux, Jean-Bernard Lubin, Jamal Green, Julia Flores, Isaiah Rozich, Tereza Duranova, Paul J. Planet, Michael A. Silverman

## Abstract

Early life microbiota drive immune system development and influence risk for immune dysfunction later in life, including the development of type 1 diabetes (T1D). Which specific early-life microbes modulate diabetes risk and the timing of these critical interactions are not well understood. To address this gap in knowledge, we screened for microbes that induce systemic IgG1 responses in young NOD mice. We isolated a strain of *Akkermansia muciniphila* that potently induces systemic IgG1 antibodies and peripheral regulatory T cells (pTregs). Since this mucus-degrading commensal protects NOD mice from T1D and is associated with lower risk of developing T1D in children, we investigated how *A. muciniphila* impacts early-life host-commensal interactions using gnotobiotic NOD mice colonized with a defined 9-member bacterial consortium that models the early life microbiome. We find that *A. muciniphila* potently induces pTregs and enhances antibody responses to other commensal microbes. Remarkably, these effects only occur when *A. muciniphila* colonizes NOD mice prior to weaning, establishing that the specific window of exposure to *A. muciniphila* shapes adaptive immune system development in diabetes-susceptible NOD mice. This time dependence provides important evidence that early-life exposure may enhance microbiota-based therapies to prevent T1D.

**One Sentence Summary:** Akkermansia muciniphila induces peripheral Tregs and enhances antibody responses to itself and other commensals during an early life window.

## Introduction

Human epidemiologic studies and experiments using animal models demonstrate that commensal microbiota profoundly influence immune system development^1^. Differences in the microbiome have been implicated in the increasing incidence of autoimmune diseases in the human population over the last half century, including increasing rates of type 1 diabetes (T1D)^2^. Specific commensal microbes are associated with increased or decreased risk of developing T1D^2–4^. It is therefore essential to identify commensals that lower risk for developing autoimmunity and understand their mechanisms of action as a critical step towards effective preventative therapies. To identify immunodulatory microbes relevant to the early stages of T1D pathogenesis, we performed a screen of intestinal commensal microbes from young non-obese diabetic (NOD) mice and identified a single microbe, *Akkermansia muciniphila*, that induces strong systemic immune responses during a critical early life period of immune system development.

*Akkermansia muciniphila* is a mucin-degrading commensal that colonizes the human gut at high abundance, comprising approximately 4% of the human fecal microbiome^5^. Higher relative abundance of *A. muciniphila* is associated with improved outcomes in a variety of immunological and metabolic disorders, including colitis^6–8^, obesity and metabolic syndromes^9–12^, as well decreasing risk of and improving treatment outcomes in intestinal cancer^13,14^. *A. muciniphila* colonization is associated with lower rates of T1D in children with elevated risk for developing this disease^4^ and reduces the incidence of T1D in NOD mice^15,16^. Yet, how *A. muciniphila* protect against the development of T1D remains poorly understood. The introduction of *A. muciniphila* to NOD mice increases the number of regulatory T cells (Tregs) in the pancreatic islets and promotes IL-10 and TGFβ expression in the islets^16^ which suggests that induction of Tregs by *A. muciniphila* may contribute to protection from developing T1D. *A. muciniphila* can also elicit antigen-specific antibody and T cell responses, including peripheral Tregs (pTregs), in C57BL/6 mice^17–19^. Identification of microbes that promote Tregs early in life before onset of autoimmunity provides high-impact therapeutic opportunity, as Tregs are key in preventing T1D in mice and humans^20–23^.

The immune system is imprinted by microbial exposures during critical windows of development in early life^24–28^. Epidemiologic studies reveal that early life antibiotics and immune exposures have long term impacts on immune function, including risk for autoimmunity, allergy and vaccine efficacy^29–31^. Studies of commensal microbiota have revealed two distinct early-life windows. The first occurs in the pre-weaning period 10 – 14 days after birth^32–35^ and the second occurs during the weaning period from 14 – 21 days after birth^25,36,37^. During the weaning period in which milk consumption declines and solid foods are introduced, there is a coordinated diversification of the microbiota and development of critical components of mucosal and systemic immunity, including pTregs^38^, mucosal IgA and systemic IgG antibodies against commensal microbes^39^. This “weaning reaction” is key for establishing homeostasis and preventing immune dysfunction later in life^24,25,34,40^. Therefore, microbes present at this early stage are poised to have a substantial impact on immune system development^24,33,40^. *A. muciniphila* colonizes the intestine of mice prior to weaning^41^, when microbial diversity is quite low^25^, suggesting that it has the potential to directly impact immune system development during these critical early-life windows.

Gnotobiotic models, which allow microbial communities to be rationally designed, are valuable tools for studying the impact of individual microbes on the host^17^ and examining the dynamics between specific microbes^42–45^. Since *A. muciniphila* colonizes well in early life when the immune system is most malleable, we introduced *A. muciniphila* into a pediatric microbiota model developed by our laboratory called Pediatric Community or PedsCom^46^. This model is comprised of 9 microbes **(Table S1)** that represent over 90% of the bacterial reads in pre-weaning mice in our specific pathogen free (SPF) NOD colony. PedsCom recapitulates the early life commensal environment in the gut and provides an ideal model in which to investigate the effect *A. muciniphila* has on cellular and humoral immune responses in early life.

Using this model, we find that *A. muciniphila* potently induces pTregs and enhances antibody responses to specific microbes in the PedsCom consortium. These effects only happen when PedsCom NOD mice are colonized with *A. muciniphila* prior to weaning. This suggests that a specific early life window of exposure is necessary for *A. muciniphila* to shape key aspects of adaptive immune development in NOD mice.

## Results and Discussion

### *A. muciniphila* induces an IgG1 antibody response in SPF NOD mice

Since CD4^+^ Tregs are protective against T1D^20,21,47^, we aimed to identify and isolate specific commensal microbes that elicit early-life diabetes-protective CD4^+^ T cell responses. Because high-throughput tools to directly identify commensal-specific CD4^+^ T cells *in vivo* are lacking, we leveraged commensal-specific systemic antibody responses to identify specific microbes that induce CD4^+^ T-cell dependent antibody responses as a way to identify microbes capable of impacting CD4^+^ T cells. We performed a screen to identify fecal microbes that elicit a mucosal IgA and/or a systemic IgG1 response in 5-week-old SPF NOD mice. We were especially interested in microbes that induce an IgG1 response since IgG1 responses to commensals are CD4^+^ T cell dependent^17^.

We leveraged an approach we call microbial flow cytometry with 16S rRNA gene sequencing (mFLOW-Seq)^48,49^, building upon IgA-seq and IgG-Seq^39,50,51^. Briefly, serum antibodies and intestinal microbes are incubated together and then microbes are sorted based on binding of IgA and IgG1 antibodies **(Fig. 1A)**. We performed the screen in 5-week-old NOD mice, the approximate age at which endogenous IgG antibodies are first produced^39^ and a time frame in which early immune infiltration of the pancreatic islets occurs^52^. We found changes in the relative abundance of bacterial classes between the antibody-negative and antibody-bound fractions; notably, there was an enrichment of Verrucomicrobiae **(Fig. 1B)**. We found a statistically significant enrichment of three microbial taxa; *Blautia*, segmented filamentous bacteria (SFB), and *Akkermansia*, were most highly enriched in the IgA and IgG1-bound fractions compared to the antibody-negative fraction **(Fig. 1C)**. When we performed the screen in 10-14 week-old NOD mice, the time period in which NOD mice typically begin to develop diabetes, *Akkermansia* was the most enriched taxa in the antibody-bound fractions (**Fig. S1A)**. We focused our studies on *A. muciniphila* because of its association with protection from T1D in mice^15,16^ and humans^4^ and its potent induction of systemic IgG1 at two key time periods important for the development of T1D.

**Figure 1.**
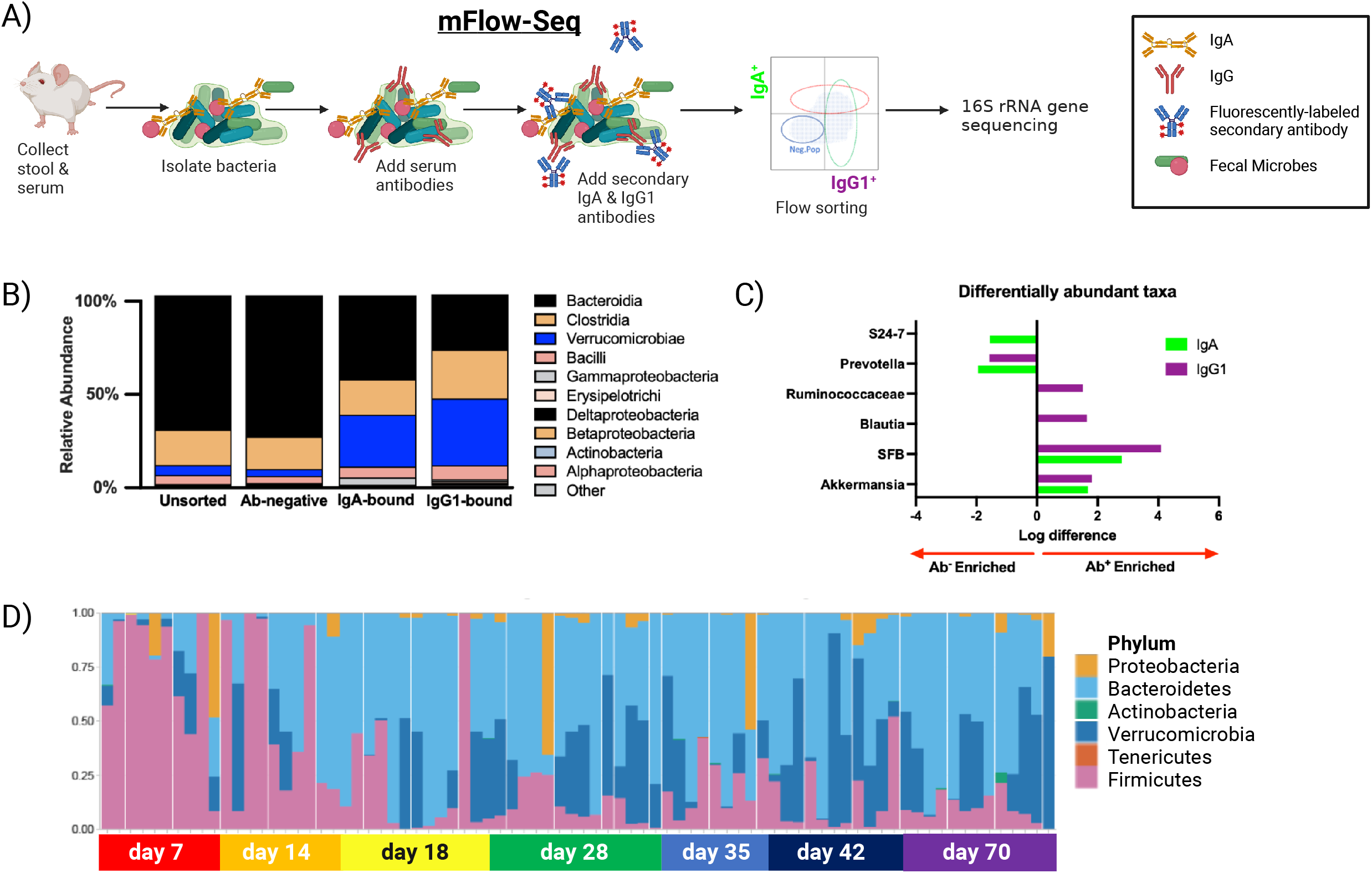
*A. muciniphila* is selectively targeted by systemic antibody responses in NOD mice. **A)** Schematic of the screening design for identifying microbes that are targeted by systemic antibodies: Stool and serum were isolated from SPF NOD mice, the bacteria were isolated from the stool and incubated with serum from the same mouse. Secondary antibodies specific for IgG1 and IgA were added, and then the bacteria were sorted based on antibody isotype bound and 16S rRNA gene sequenced. **B)** Relative abundance of bacterial classes present in the unsorted, antibody-negative, and IgA and IgG1 bound fractions of 5-week-old NOD mice (n = 8). **C)** Bacterial taxa with the highest differential abundance in the antibody-bound fraction compared to the antibody-negative fraction from the mice in (B) as determined by a linear mixed-effects model (FDR, significance <0.1). **D)** Relative abundance of bacterial phyla in stool from SPF NOD mice over the first 10 weeks of life.

Since *Akkermansia* is present in the fecal microbiomes of children^4^ and NOD mice^15^, we analyzed *A. muciniphila* colonization in our SPF NOD colony from birth through 10 weeks of age to determine when this commensal microbe colonizes NOD mice. We found that *A. muciniphila* in fecal samples was present at low levels at birth, expanded at day of life 7 and was dynamic throughout the life of individual mice **(Fig. S1B)**; it varied within litters, and was often present in high abundance in pre-weaning NOD mice. Specifically, high levels of *A. muciniphila* were present during the early-life pre-weaning (days of life 0 – 14) and weaning period (days of life 14 – 21) **(Fig. 1D)**. We isolated and cultured a strain of *A. muciniphila* present in our SPF NOD colony and confirmed its taxonomy by sequence analysis of the complete 16S rRNA gene and whole genome (manuscript in preparation). Thus, early life colonization positions *A. muciniphila* to be a potent immune modulator, as this period is key in immune system development and pathogenesis of autoimmune diabetes^53^.

### *A. muciniphila* induces RORγ^+^ Tregs in NOD mice in early life

Since *A. muciniphila* colonizes NOD mice during a period of rapid immune system development, we hypothesized that pre-weaning colonization would impact adaptive immune responses to a greater extent than exposure to *A. muciniphila* later in life. To achieve natural acquisition, we used this strain of *A. muciniphila* isolated from our SPF NOD mice to colonize germfree (GF) NOD dams prior to pregnancy to investigate the effect of vertical transfer of *A. muciniphila* on the resulting offspring. We focused on the development of RORγ^+^Helios^-^Foxp3^+^ peripheral Tregs (pTregs) **(Fig. S2A)**, a subset of Tregs that develops around weaning in response to changes in commensal microbes and microbial metabolites^25,38^. Peripheral Tregs are relevant to the pathogenesis of T1D, as they restrain insulitis and T1D in NOD mice^54^. We found that vertical transfer of *A. muciniphila* to GF NOD mice leads to increased proportions of Foxp3^+^ Tregs in the lamina propria of the cecum and colon **(Fig. 2A-B)** and increased proportions of RORγ^+^Foxp3^+^ Tregs in the cecal and colonic lamina propria as well as the spleen, mesenteric lymph nodes (MLNs), and pancreatic lymph nodes (PLNs) **(Fig. 2C-D)**.

**Figure 2.**
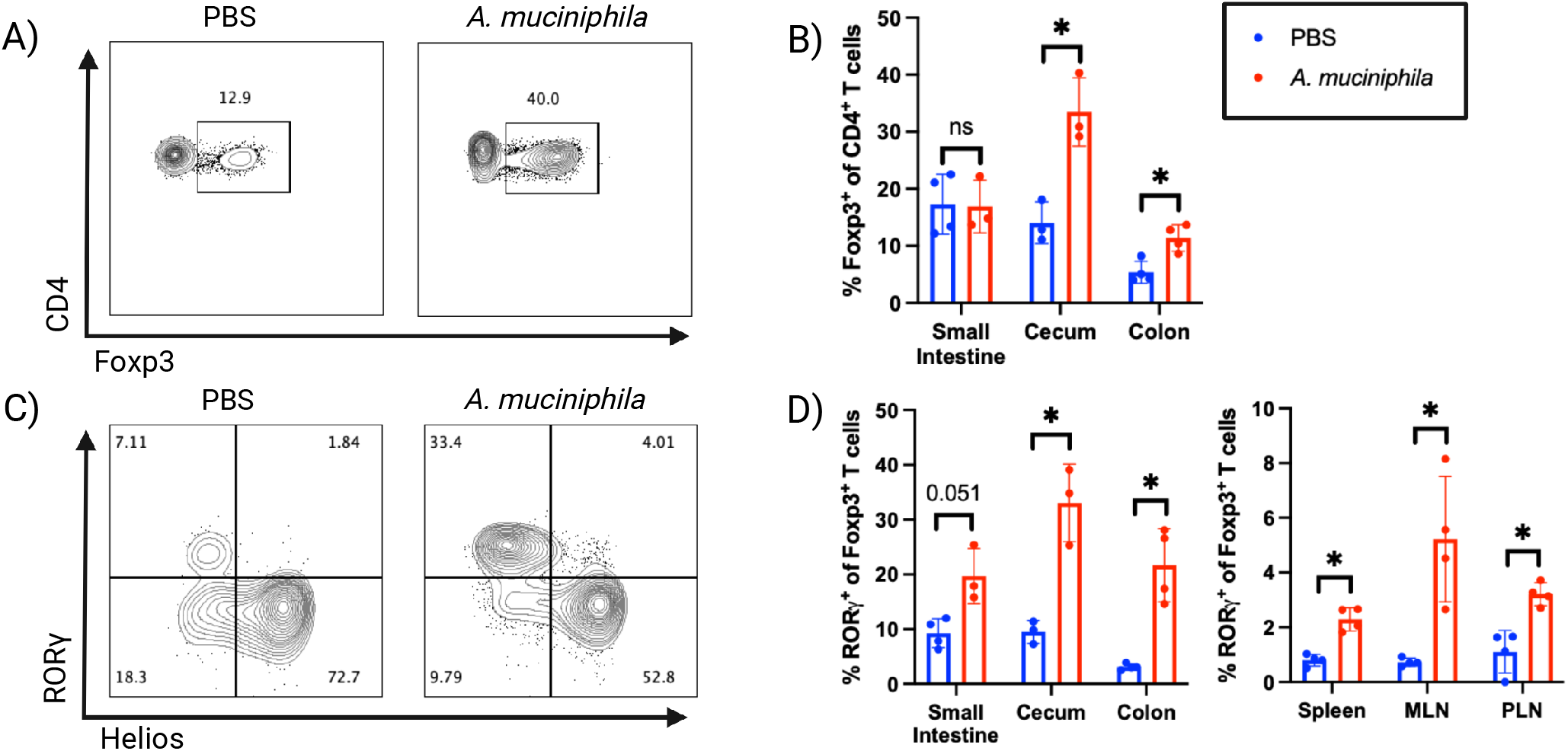
*A. muciniphila* increases Foxp3^+^ Tregs and RORγ^+^ Tregs in germ free NOD mice. Representative flow plots showing Foxp3^+^ Tregs **(A)** and RORγ^+^Foxp3^+^ Tregs **(C)** in the cecum of GF NOD mice born to dams gavaged with PBS or 1 × 10^9^ CFU *A. muciniphila* prior to pregnancy. Percentage of Foxp3^+^ Tregs **(B)** in the lamina propria of GF NOD mice and percentage of RORγ^+^Foxp3^+^ Tregs **(D)** in the lamina propria and lymphoid tissues of GF NOD mice in (A-B). Data are from 1 experiment, represented as mean +/- standard deviation (SD). Statistical significance was determined using the Mann-Whitney test, *p-value < 0.05, **p-value < 0.01, ***p-value < 0.001.

To study its effects in the context of the early life gut, we introduced *A. muciniphila* into a gnotobiotic model of the early life microbiota called PedsCom, which is a tractable model designed to study the impact of specific commensal microbes on early life immune system development. Unexpectedly, introducing *A. muciniphila* at weaning **(Fig. 3A)** did not have any discernable effect on proportions of RORγ^+^Foxp3^+^ Tregs **(Fig. 3B and Fig. S2C)**. Since we saw a dramatic increase in pTregs following vertical transmission of the monocolonized NOD mice, we hypothesized that *A. muciniphila* may impact immune system development during the pre-weaning period. Indeed, when PedsCom NOD neonates were colonized by vertical transfer from dams gavaged with *A. muciniphila* prior to pregnancy **(Fig. 3A)**, we found striking increases in the proportions and total numbers of RORγ^+^Foxp3^+^ Tregs in the spleen, MLNs, and lamina propria of the cecum and colon of the offspring **(Fig. 3B-C and Fig. S2B)**. These data suggest that pTreg induction by this strain of *A. muciniphila* is restricted to a pre-weaning life window in context of PedsCom NOD mice.

**Figure 3.**
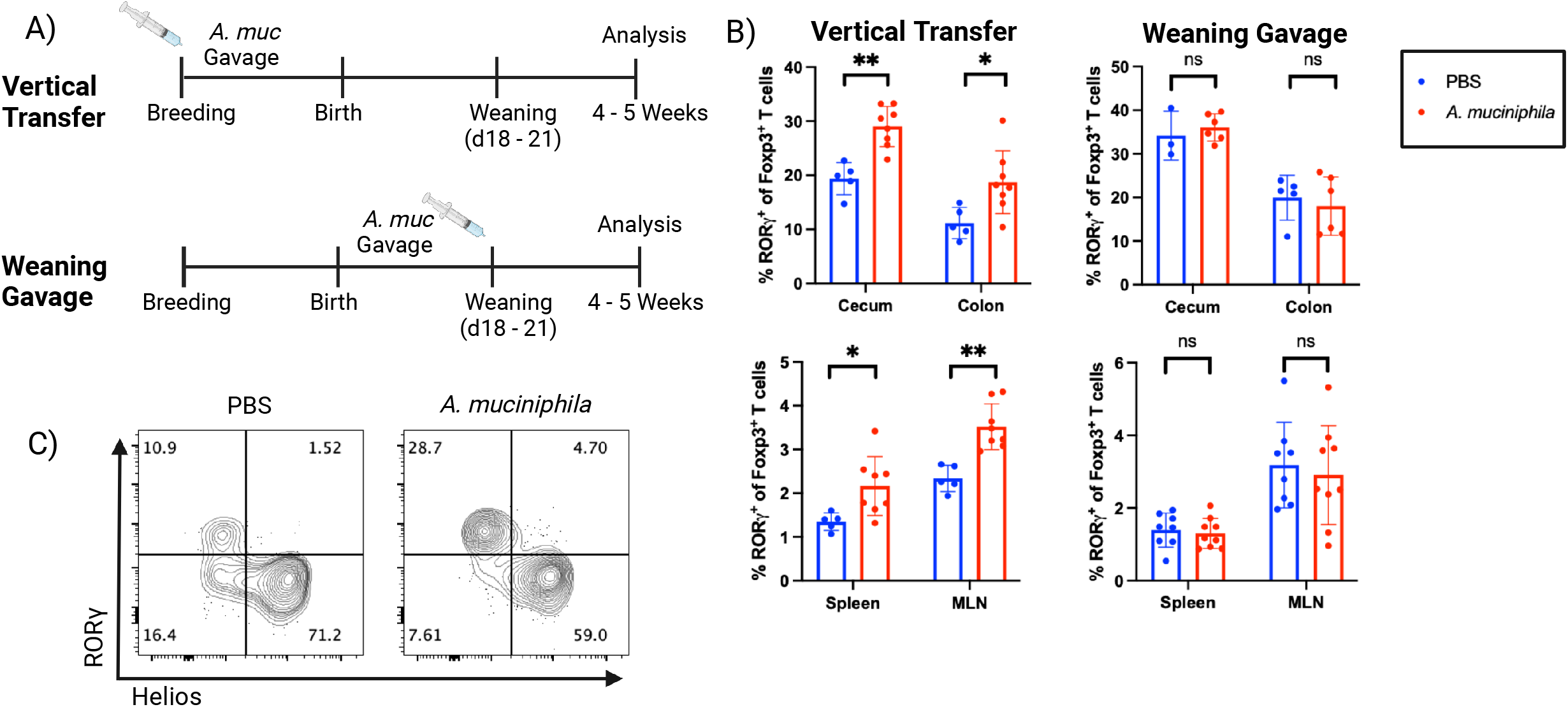
*A. muciniphila* induces RORγ^+^ Tregs in pre-weaning but not weaning age PedsCom NOD mice. **A)** Schematic showing the timeline of *A. muciniphila* colonization and analysis of RORγ^+^Foxp3^+^ Tregs in Pedscom colonized NOD mice. In the vertical transfer model (top), dams were gavaged with PBS or 2 × 10^9^ CFU *A. muciniphila* prior to or during pregnancy and Tregs were analyzed in the offspring at 4 – 5 weeks of age. In the weaning age model (bottom), PedsCom colonized mice were gavaged with PBS or 2 × 10^9^ CFU *A. muciniphila* at days 18 – 20 and analyzed for Treg induction at 5 weeks of age. **B)** RORγ^+^Foxp3^+^ Tregs as a percentage of Foxp3^+^ Tregs in the cecal and colonic lamina propria, spleen, and MLNs of PedsCom colonized mice colonized by vertical transfer (left) and analyzed at 4-5 weeks or gavaged directly with *A. muciniphila* at weaning and analyzed at 5 weeks of age (right). Data are representative of and pooled from 2 experiments, respectively. Data are represented as mean +/- SD. Statistical significance was determined using the Mann-Whitney test, *p-value < 0.05, **p-value < 0.01, ***p-value < 0.001. **C)** Representative flow plots of RORγ^+^Foxp3^+^ Tregs in the ceca of 4-week-old PedsCom colonized NOD mice born to dams gavaged with PBS or *A. muciniphila* (vertical transfer).

### *A. muciniphila* enhances systemic antibody responses to staphylococcal species in young PedsCom NOD mice

Due to its close proximity to the intestinal epithelium, we hypothesized that *A. muciniphila* would stimulate strong mucosal and systemic antibody responses. As expected, we found that vertical transfer of *A. muciniphila* in PedsCom NOD mice induced strong systemic IgA, IgG1, and IgG2b responses against *A. muciniphila* (**Fig. 4A-B)**. Colonizing adult PedsCom NOD mice with *A. muciniphila* induced systemic IgA and IgG2b responses against *A. muciniphila* but lead to a minimal IgG1 response **(Fig. 4A, C)**. Since IgG1 responses against commensal microbes are CD4^+^ T cell-dependent^17^, these findings further supports a model in which *A. muciniphila* selectively induces T cell responses before weaning.

**Figure 4.**
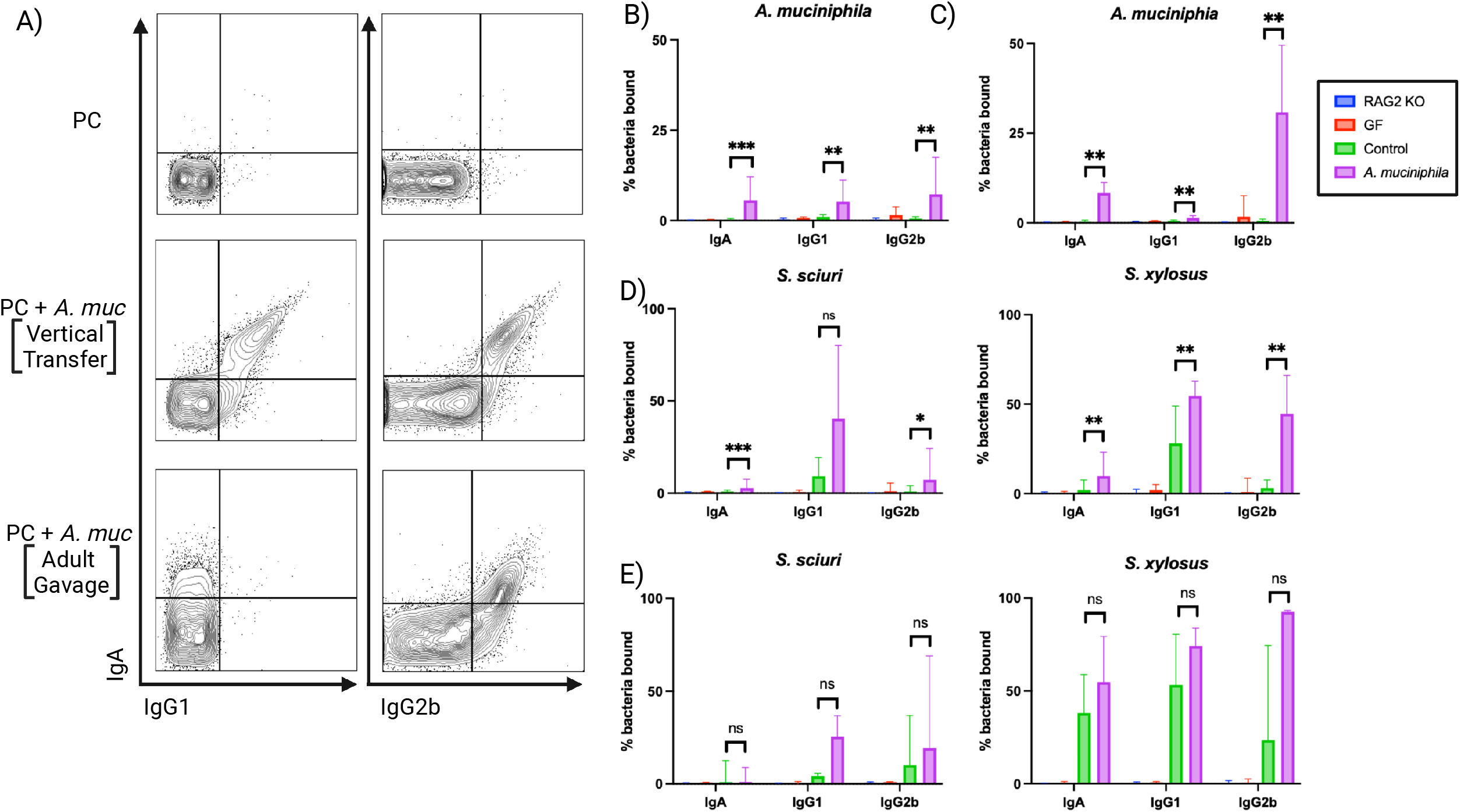
Vertical transfer of *A. muciniphila* enhances antibody response to PedsCom staphylococcal species. **A)** Representative flow plots showing serum antibody binding to cultured *A. muciniphila* from control 6-week-old PedsCom NOD mice (top), 6-week-old PedsCom NOD mice born to dams gavaged with 2 × 10^9^ CFU *A. muciniphila* prior to pregnancy (middle), and 10 to 12-week-old PedsCom NOD mice gavaged with 2 × 10^9^ CFU *A. muciniphila* 2 weeks prior to analysis (bottom). Quantification of median binding of serum from control mice and mice born to *A. muciniphila*-colonized dams **(B)** and adult mice gavaged with PBS or *A. muciniphila* **(C)** from (A), as measured by mFLOW. **D)** Median binding to *S. sciuri* and *S. xylosus* of serum antibodies from the mice in (A) that were colonized with *A*. muciniphila by vertical transfer, measured by mFLOW. **E)** Binding to *S. sciuri* and *S. xylosus* of serum antibodies from 10 to 12-week-old mice colonized by 2 × 10^9^ CFU *A. muciniphila* two weeks prior to analysis. Data are pooled from 2 – 4 independent experiments. Data are represented as median +/- interquartile range. Statistical significance was determined using the Mann-Whitney test, *p-value < 0.05, **p-value < 0.01, ***p-value < 0.001.

Since *A. muciniphila* has cooperative^55^ or antagonistic^42^ relationships with other commensals that influence interactions with the host, we next leveraged the PedsCom mice to determine if *A. muciniphila* colonization shapes host response to other microbes. Using microbial flow cytometry, we compared the antibody responses of PedsCom NOD mice with or without vertical colonization of *A. muciniphila*. We found enhanced binding of systemic IgA and IgG2b to *Staphyloccus sciuri* and systemic IgA, IgG1, and IgG2b to *Staphyloccus xylosus*, the two staphylococcal species in PedsCom (**Fig. 4D** and **Table S1**). This enhanced systemic antibody response is specific to the staphylococcal species as there was no enhancement of antibody responses to the other seven PedsCom microbes (**Fig. S3**). In contrast to the preweaning colonization, PedsCom NOD mice colonized with *A. muciniphila* during adulthood did not demonstrate enhanced antibody responses to any of the PedsCom microbes (**Fig. 4E**), emphasizing the importance of early-life exposure to *A. muciniphila* for enhanced systemic antibody responses to other commensal microbes. This observation is not explained by cross-reactivity between *A. muciniphila*-specific antibodies and staphylococcal epitopes, as vertical colonization of GF NOD mice with *A. muciniphila* did not result in an increase in antibody binding to *S. sciuri*, although we did observe a modest increase in IgG1 binding to *S. xylosus* (**Fig. S4**). These data support a model in which *A. muciniphila* colonizes the intestinal mucosa of mice during a pre-weaning window of development and stimulates systemic antibodies and Treg responses to itself and potentially other microbes.

Vertical transfer of *A. muciniphila* in PedsCom C57BL/6 mice did not result in an increase in RORγ^+^Foxp3^+^ Tregs (**Fig. S5**) or enhancement of antibody binding to PedsCom microbes **(Fig. S6)**, indicating that diabetes-susceptible NOD mice have distinct responses to *A. muciniphila* in the context of PedsCom consortium of microbes.

In this study, we identified *A. muciniphila* as a strong inducer of adaptive immune responses in NOD mice, including systemic IgG and RORγ^+^ pTregs in gut and lymphoid tissues. Importantly, *A. muciniphila* was associated with lower risk of progression to T1D in the TEDDY trial^4^ and murine studies implicated this microbe in decreasing T1D rates in NOD mice^15,16^; however, the specific mechanisms by which it functions in NOD mice remain poorly understood. We determined that the ability of *A. muciniphila* to enhance RORγ^+^Foxp3^+^ Treg induction and commensal-targeting systemic antibody production are age restricted in NOD mice, with preweaning exposure to *A. muciniphila* being necessary for the microbe to modulate these immune responses in the context of a defined set of 9 bacteria that models the early-life microbiome^46^. Additionally, we demonstrated that the presence of *A. muciniphila* enhances antibody responses to Staphylococcus species in NOD mice colonized with PedsCom. This finding suggests a novel mechanism by which *A. muciniphila* shapes adaptive immune responses by altering immune responsiveness to other immunostimulatory commensal microbes.

Age at time of colonization is a key host factor that affects immune responses to *A. muciniphila*. In this study, we find that colonizing PedsCom mice with *A. muciniphila* through natural acquisition at birth induces RORγ^+^Foxp3^+^ Tregs and enhances antibody responses to other commensals. Specifically, we find that colonization before weaning is important for expansion of pTregs in intestinal and lymphatic tissues. A similar requirement for pre-weaning colonization has been described restraining intestinal natural killer T cell (iNKT) development by the human commensal microbe *B. fragilis*^35^. To our knowledge, our data provide the first example of a specific commensal microbe derive from mice impacting adaptive immune response in a strictly early life context in a model of autoimmunity.

By colonizing early in life, we propose that *A. muciniphila* is poised to shape the critical immunologic development that occurs during the preweaning and weaning period. This weaning period includes opening of GAPs in the colon which facilitates development of commensal specific pTregs^36,37^ and anti-commensal IgA and IgG antibodies. This is a life stage that coincides with a substantial diversification of the microbiota and extensive immune development, a phenomenon called the “weaning reaction”^25^. This environment is especially amenable to developing tolerogenic CD4^+^ T cell responses to microbial signals^32^. Our data showing that Treg and antibody induction by *A. muciniphila* is restricted to this early time frame in NOD mice indicates that *A. muciniphila* may promote homeostasis early in life and interact with the immune system less during adulthood. This strong dependence on early-life colonization for the immunomodulation suggests that designing effective *A. muciniphila* based therapeutics to prevent T1D may require targeting specific early life developmental window.

*A. muciniphila* directly impact the host immune system by a variety of well-described mechanisms including inducing antigen-specific CD4^+^ T cells^17–19^ and systemic IgG antibodies^17^, and stimulating innate immunity via toll-like receptor agonists^56–58^. Here we provide evidence that early-life exposure to *A. muciniphila* can also impact the immune system by enhancing immune responses to other commensal microbes. *A. muciniphila* enhances antibody responses specifically to the PedsCom staphylococcal species *S. sciuri* and *S. xylosus* when mice are colonized at birth but not after colonization during adulthood. The initiation or enhancement of antibody responses to specific commensal bacteria by another commensal microbe has not been previously reported to our knowledge, as similar effects have only been observed in the context of pathogenic infections like *Helicobacter bilis*^59^. In a pathogenic context, these “off-target” immune responses are generally attributed to breach of the intestinal barrier and increased translocation of other microbes or microbial antigens in the context of inflammation. Since *A. muciniphila* is generally considered to be anti-inflammatory, this effect could represent a novel mechanism by which commensals may promote tolerance to the microbiota in early life. Future studies are needed to determine how *A. muciniphila* enhances commensal antibody targeting of these two staphylococcal species. *A. muciniphila* has been shown to enhance barrier function, but its effect on immune detection of and response to gut antigens has yet to be determined. *A. muciniphila* is known to promote goblet cell hyperplasia, so it is possible it could enhance antigen passage through increased goblet cell associated antigen passage (GAP) formation during early life^36^. Further study using gnotobiotic mice may help unravel the degree to which direct or indirect effects of *A. muciniphila* are driving protection from developing autoimmunity.

One limitation of this study is that it does not establish the TCR specificity of the RORγ^+^Foxp3^+^ pTregs induced in the NOD mice. Based on previous studies in C57BL/6 mice, we anticipate that some of the pTregs are specific for *A. muciniphila* antigens^17^. However, given the changes in antibody responses to other Staphylococcus species, it is also possible that some of these microbially-induced pTregs may be specific for antigens from other commensal microbes. In addition to determining specificity, it will also be important to determine the effect of these pTregs on T1D pathogenesis.

*A. muciniphila* is associated with better outcomes in a variety of disease states including T1D, but the mechanism by which it prevents autoimmunity is unclear. Here, we establish that early-life exposure is necessary for *A. muciniphila* to induce RORγ^+^Foxp3^+^ Tregs and enhance commensal-targeting antibody responses. This discovery provides a better understanding of how and when *A. muciniphila* interacts with other commensals in a genetically T1D-prone host, and provides support for focusing on the early-life period to design microbiota-based therapies to prevent T1D. This work also highlights the utility of the designed gnotobiotic model PedsCom for dissecting the effects of individual immunomodulatory microbes in early life.

## Methods and materials

### Mice

Specific pathogen free NOD were bred and housed in the Colket Translational Research Building (CTRB) vivarium at the Children’s Hospital of Philadelphia. CTRB-housed mouse colonies were fed Lab Diet 5058 (Cat# 0001329) throughout maintenance and experiments. Germ-free and gnotobiotic C57BL/6J and NOD mice were sterilely derived and kept in plastic isolators [Class Biologically Clean (CBClean), WI, USA] in the Hill Pavilion gnotobiotic core at the University of Pennsylvania. They were fed autoclaved sterile Lab Diet 5021 (Cat# 0006540) and housed on autoclaved, sterile Beta-chip hardwood bedding (Nepco, NY, USA). The isolators were checked for bacterial contamination monthly by plating fecal pellets on brain heart infusion (BHI) (Oxoid, UK), NB1, and Sabouraud media agar at 37°C for 65-70 hours aerobically and anaerobically with negative and positive controls. The isolators were independently tested for contamination quarterly by Charles River Laboratories (NJ, USA). PedsCom NOD and C57BL/6J mice were generated as previously described^46^. Gnotobiotic mice were transferred sterilely to autoclaved cages containing the same bedding and food as isolator cages and housed outside isolators during experiments that required colonizing with *A. muciniphila*. Mice were handled sterilely throughout experiments and fecal pellets were collected weekly or biweekly and analyzed by qPCR to ensure no contamination of control groups with *A. muciniphila*.

### Bacterial Culturing

PedsCom bacteria were cultured on BHI agar plates supplemented with 1% BBL vitamin K1-Hemin solution (BHI-KH) aerobically or under anaerobic conditions (90% N2, 5% CO2, 5% H2) at 37°C. BHI + KH media was inoculated and cultured overnight or for 48 hours for slower growing species (*P. distasonis). A. muciniphila* was isolated from fecal pellets from an 18-day-old NOD mouse in CTRB and cultured on minimal media^60^ + 0.1% porcine stomach mucin (MMM) agar plates followed by liquid culture in MMM overnight. Colonies were screened using primers specific for the Verrucomicrobiota 16S gene (primer sequences-Forward: TCA KGT CAG TAT GGC CCT TAT, Reverse: CAG TTT TYA GGA TTT CCT CCGCC). One confirmed isolate was streaked on MMM and BHI-KH + 0.1% mucin (BHI-KHM) and was sent for 16S rRNA gene metagenomic sequencing and whole genome sequencing and then frozen in 25% glycerol at -80°C to serve as a bacterial stock. For gavaging and mFLOW experiments, *A. muciniphila* was streaked from those stocks on BHI-KHM agar plates and grown overnight in liquid BHI-KHM cultures. Those cultures were centrifuged at 4°C at 4000 x g, and resuspended in sterile, reduced 0.1% L-cysteine, phosphate buffered saline (rPBS). The optical density (OD) of the culture was determined using a spectrophotometer and the bacterial concentration adjusted to an OD600 of 0.1 or 0.01 for mFLOW experiments and to 2 × 10^10^ CFU/mL for gavage experiments.

### Bacterial Inoculation

NOD or C57BL/6 mice 6-10 weeks of age were orally gavaged with 100 µL rPBS or 2 × 10^9^ CFU *A. muciniphila* in 100 µL rPBS using a sterile syringe and a straight 1-inch-long 22-gauge gavage needle (Cadence Science, Cat#7901). Mice younger than 6 weeks were gavaged using 1 inch long, 24-gauge needle and 50 µL rPBS or 1 × 10^9^ CFU *A. muciniphila* in 50 µL rPBS.

### Colonization Monitoring

Fecal pellets or intestinal contents were collected in sterile microcentrifuge tubes and stored at -80°C. Bacterial DNA was isolated from samples using the DNAeasy PowerSoil kit (Qiagen, Germany) using a QIACube (Qiagen, #9001292). *A. muciniphila* colonization was determined by qPCR, using PowerUp SYBR Green Master Mix (Thermofisher cat# A25741) and primers for the *rpoB* gene of *A. muciniphila* (primer sequences – Forward: GTTCACGACCACATCGAAA, Reverse: CGTCATCCAGTTCCGTAAAG).

### Microbial flow cytometry (mFLOW)

Fecal pellets were homogenized in 1 mL PBS, 1% BSA (PBS-B) and centrifuged at 100 x g for 15 minutes to pellet solid material. Fecal supernatants were transferred to sterile microcentrifuge tubes and centrifuged at 8000 x g for five minutes to pellet the bacterial fraction and resuspended to an OD600 of 0.1 in PBS-B. Cultured bacteria were centrifuged at 4700 x g for 10 minutes, resuspended in 1 mL PBS, 1% BSA, and diluted to an OD600 of 0.1 or 0.01. Bacteria were washed with PBS-B, resuspended in blocking buffer (PBS-B, 20% normal rat serum), and incubated for 30 minutes at 4°C. The bacteria were then incubated with serum (inactivated at 56°C for 30 min, centrifuged at 16000 x g for 5 min at 4°C) diluted 1:25 in PBS, 1% BSA, sodium azide (BSB) at 4°C for 1 hr. The bound bacteria were then washed with BSB and incubated with anti-mouse antibodies (PE-IgA, clone mA-6E1; PE-Cy7-IgG1, clone RMG1-1; BV711-IgG2b, clone R12-3; BV421-IgG3, clone R40-82) diluted 1:25 in BSB for 30 minutes at 4°C, washed and incubated in SytoBC nucleic acid stain (Invitrogen #S34855) diluted 1:500 in Tris-buffered saline (TBS) for 15 minutes at room temperature. Flow cytometry analysis on stained bacteria populations with SytoBC and isotype-stained controls was used to define living bacteria. Antibody-bound bacteria were sorted on a FACSAria Fusion Sorter and the antibody-bound and unbound fractions submitted for 16S rRNA gene sequencing.

### 16S rRNA gene metagenomic sequencing

The PennCHOP microbiome program sequencing core sequenced the V4 variable region of the 16S rRNA gene using the Illumina MiSeq platform as previously described^61^. QIIME2 (ver. 2018.2) was used to analyze the resulting sequencing libraries^62^ and DADA2 was used to cluster and de-noise the results into amplicon sequence variants. The greengenes 99% rRNA gene reference database (ver. 13.8) was used to classify bacterial taxons. Taxa bar plots were generated with the R package phyloseq^63^.

### Isolation & characterization of lamina propria and lymphoid T cells

Mouse spleen, MLN, PLN, small intestine (SI), cecum, and colon tissues were isolated from euthanized mice and placed in cell culture media (CCM: RPMI, 100 units/mL penicillin, 100 µg/mL streptomycin, 10% fetal bovine serum (FBS)). The lymph nodes and spleens were dissociated using the back of the plunger of a 1 mL syringe through a 70 µM filter in 1 mL CCM in a dish. The dish and filter were rinsed with CCM and the suspension was spun down at 480 x g for 10 minutes in 15 mL conical tubes. The supernatant was discarded, and MLN and PLN pellets were resuspended in 200 µL CCM and plated in a 96 well round bottom plates for staining. Splenocytes were resuspended in 1 mL ACK lysis buffer for 1 minute; they were then quenched with 10 mL CCM and centrifuged as before. The spleen pellet was then resuspended in 1 mL staining media (RPMI, 4% FBS) and 100 µL of each sample was plated for staining. The intestinal tissues were cleared of debris by gently flushing with PBS using a gavage needle (SI & colon), or by cutting open and shaking in CCM (cecum). The cecal patch and Peyer’s patches from the small intestine were excised. The SI and colon were rolled on a paper towel and a scalpel was used to remove mesenchyme. All tissues were then cut open and shaken to remove remaining intestinal debris. The intestinal tissues were then incubated in intraepithelial lymphocyte (IEL) digestion media (30 mL RPMI, 0.062% dithiothreitol, 1 µM EDTA, 1.3% FBS) and incubated while stirring at 300 rpm with a magnetic stir bar at 37°C for 15 minutes to remove IEL cells. The tissues were washed with PBS, placed in plain RPMI, and diced into small sections using dissection scissors. The tissues were then digested for 40 minutes in 25 mL digestion media (25 mL RPMI, 12.5 mg dispase (Gibco), 37.5 mg collagenase type II (Gibco), 1 mL PBS) with stirring at 300 rpm at 37°C to create a single cell suspension. The suspension was filtered through 100 µM filters and quenched with 25 mL RPMI. The solutions were centrifuged for 10 minutes at 310 x g at 4°C, resuspended in approximately 200 µL staining media and plated for staining. Staining for surface markers of T cells was carried out in the dark at 4°C for 15 – 30 minutes using the following antibody panel: BV510-CD45 (clone 30F11; Biolegend, CA, USA), PE-Cy7-TCRβ (clone H57-597; Biolegend), APC-Cy7-CD19 (clone 6D5; Biolegend), AF700-CD8 (clone 53-6.7; Biolegend), FITC-CD4 (clone Rm4-5; Biolegend). Cells were washed 2X with staining media and fixed overnight at 4°C using the eBioscience Fixation kit (cat# 005223-56, 00-5123-43). Cells were washed 2X in permeabilization buffer (Invitrogen cat# 00-8333-56) with centrifugation at 1340 x g and stained for intracellular proteins for 50 minutes at room temperature using the following panel: APC-Foxp3 (FJK-16s; eBioscience, MA, USA), PE-RORγ (clone B2D; eBioscience), Pacific Blue-Helios (clone 22F6; Biolegend). Cells were washed 2X with permeabilization buffer, resuspended in 200 µL PBS, and filtered through 50 µM nylon mesh (Genesee Scientific, cat# 57-106) into FACS tubes. Stained samples were analyzed on the LSRFortessa (BD) and data analysis was performed on FlowJo v10 software (BD).

### Bioinformatics and statistics

Statistical analyses were performed using non-parametric Mann Whitney-Wilcoxon tests on median values with Prism 8 (GraphPad Software Inc), except where noted otherwise. P values <0.05 were considered statistically significant. Differential abundance analysis (Fig. 1C and Fig. S1A) was performed on logit transformed relative abundance of taxa greater than 1% across all samples. A linear mixed-effects model was performed on transformed data with population and genotype as fixed effects and cage as the random effect. Multiple tests were corrected for by FDR, significance <0.1.

## Acknowledgements

The authors thank Dr. Joseph Zackular, Dr. Paul Planet, Dr. Lawrence Eisenlohr, Dr. Paula Oliver, Dr. Jorge Henao-Mejia, Dr. Maayan Levy, and Dr. Taku Kambayashi for helpful input. The authors also thank Dmitri Kobuley and Michelle Albright of the PennCHOP microbiome program gnotobiotic mouse facility at Hill Pavilion. Figures made using BioRender.

## Author Contributions

Conceptualization, S.M. and M.A.S.; Methodology, S.M. and M.A.S..; Investigation, S.M., JL, and M.A.S..; Writing – Original Draft, S.M. and M.A.S..; Writing – Review & Editing, S.M. and M.A.S..; Funding Acquisition, S.M. and M.A.S.; Resources, M.A.S.; Supervision, M.A.S.

## Supplemental Figure Legends

**Figure S1. *A. muciniphila* colonization is dynamic throughout life and is enriched for antibody binding in 10-week-old SPF NOD mice. A)** Bacterial taxa with the highest differential abundance in the antibody-bound fraction compared to the unbound fraction from 10-week-old SPF NOD mice as determined by a linear mixed-effects model (FDR, significance <0.1). **B)** *A. muciniphila* colonization of individual SPF NOD mice over the first 10 weeks of life, measured by qPCR.

**Figure S2. RORγ**^**+**^, **Helios**^**-**^, **Foxp3**^**+**^ **Tregs: gating strategy and changes in numbers in PedsCom NOD mice. A)** Gating strategy for identifying RORγ^+^Foxp3^+^ Tregs in the lamina propria (CD45^+^, singlet, TCRβ^+^, CD4^+^, Foxp3^+^, RORγ^+^, Helios^-^). Results from a cecal sample used as an example. **B)** Numbers of RORγ^+^Foxp3^+^ Tregs in the cecal and colon lamina propria, spleen, and MLN of 4-week-old PedsCom colonized NOD mice born to dams gavaged with PBS or *A. muciniphila* (vertical transfer, left) or PedsCom colonized NOD mice gavaged directly with *A. muciniphila* at weaning and analyzed at 5 weeks of age (weaning gavage, right). Data representative of and pooled from 2 experiments, respectively. Data are represented as mean +/- SD. Statistical significance was determined using the Mann-Whitney test, *p-value < 0.05, **p-value < 0.01, ***p-value < 0.001.

**Figure S3. Vertically transferred *A. muciniphila* does not alter antibody responses to most species in PedsCom NOD mice**. Binding of serum antibodies to cultured PedsCom bacteria from 6-week-old PedsCom NOD mice born to dams gavaged with PBS or 2 × 10^9^ CFU *A. muciniphila* prior to pregnancy. Data pooled from four experiments, represented as median +/- interquartile range. Statistical significance determined using the Mann-Whitney test, *p-value < 0.05, **p-value < 0.01, ***p-value < 0.001.

**Figure S4. Antigen cross-reactivity does not fully explain *A. muciniphila* mediated enhancement of antibody responses to PedsCom staphylococcal species**. Binding of serum antibodies to some cultured PedsCom bacteria from GF NOD mice born to dams gavaged with PBS or 2 × 10^9^ CFU *A. muciniphila* prior to pregnancy. Data are from 1 experiment, represented as mean +/- SD. Statistical significance was determined using the Mann-Whitney test, *p-value < 0.05, **p-value < 0.01, ***p-value < 0.001.

**Figure S5. Vertically transferred *A. muciniphila* does not induce RORγ**^**+**^ **Tregs in PedsCom C57BL/6 mice. A)** Foxp3^+^ Tregs as a percentage of CD4^+^ T cells in the indicated tissues from 6-week-old PedsCom NOD mice born to dams gavaged with PBS or 2 × 10^9^ CFU *A. muciniphila* prior to pregnancy. **B)** RORγ^+^Foxp3^+^ Tregs as a percentage of Foxp3^+^ Tregs in the lamina propria and lymphoid tissue of the mice in A). Data are from 1 experiment, represented as mean +/- SD. Statistical significance was determined using the Mann-Whitney test, *p-value < 0.05, **p-value < 0.01, ***p-value < 0.001.

**Figure S6. Vertically transferred *A. muciniphila* does not alter antibody responses to PedsCom microbes in C57BL/6 mice**. Binding of serum antibodies to cultured PedsCom bacteria from 6-week-old PedsCom C57BL/6 mice born to dams gavaged with PBS or 2 × 10^9^ CFU *A. muciniphila* prior to pregnancy. Data are pooled from four experiments, represented as median +/- interquartile range. Statistical significance was determined using the Mann-Whitney test, *p-value < 0.05, **p-value < 0.01, ***p-value < 0.001

