## Supplemental Table 1 for "Modulation of adaptive immune responses by *Akkermansia muciniphila* is restricted to an early life window in NOD mice"

### Table S1. Bacterial species in the PedsCom consortium

| **Genus and Species** | **Phylum** | **Family** | **Genome size (MB)** |
|---|---|---|---|
| *Staphylococcus xylosus* | Firmicutes | Staphylococcaceae | 2.83 |
| *Staphylococcus sciuri* | | | 2.84 |
| *Enterococcus faecalis* | | Enterococcaceae | 2.88 |
| *Lactobacillus murinus* | | Lactobacillaceae | 2.48 |
| *Lactobacillus johnsonii* | | | 1.92 |
| *Anaerostipes caccae* | | Lachnospiraceae | 3.39 |
| *Clostridium intestinale* | | Clostridiaceae | 4.69 |
| *Kosakonia cowanii* | Proteobacteria | Enterobacteriaceae | 4.99 |
| *Parabacteroides distasonis* | Bacteroidetes | Tannerellaceae | 5.33 |
