## Supplementary Figures for "Modulation of adaptive immune responses by *Akkermansia muciniphila* is restricted to an early life window in NOD mice"

Figure S1. *A. muciniphila* colonization is dynamic throughout life and is enriched for antibody binding in 10-week-old SPF NOD mice

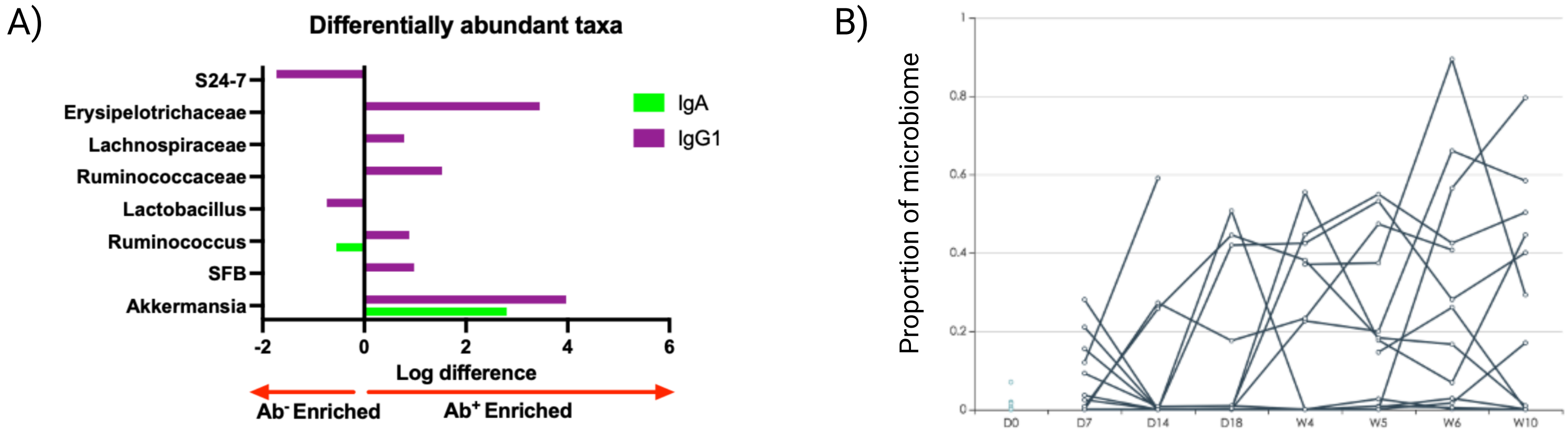

Figure S2. RORγ⁺, Helios⁻, Foxp3⁺ Tregs: gating strategy and changes in numbers in PedsCom NOD mice

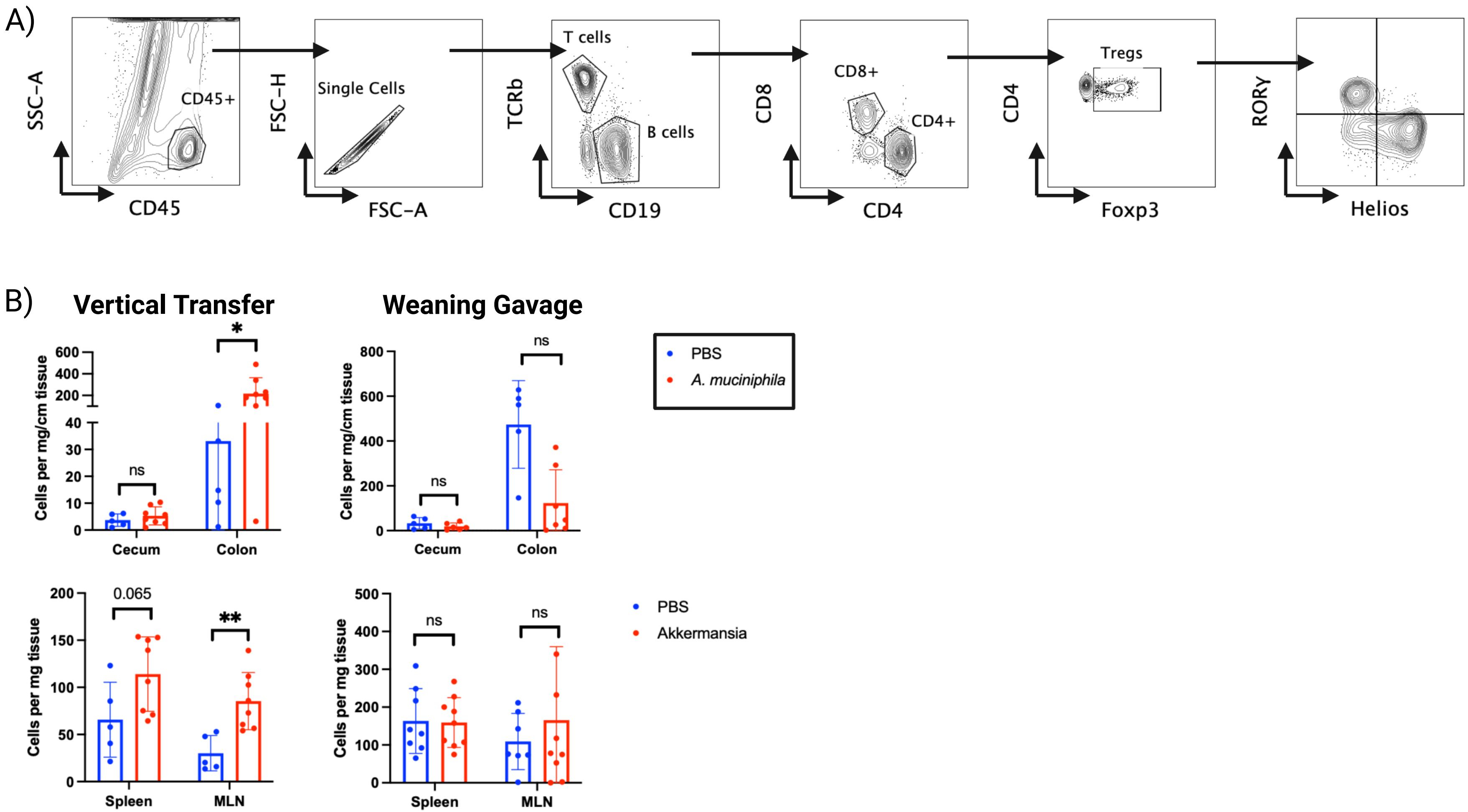

Figure S3. Vertically transferred *A. muciniphila* does not alter antibody responses to most species in PedsCom NOD mice.

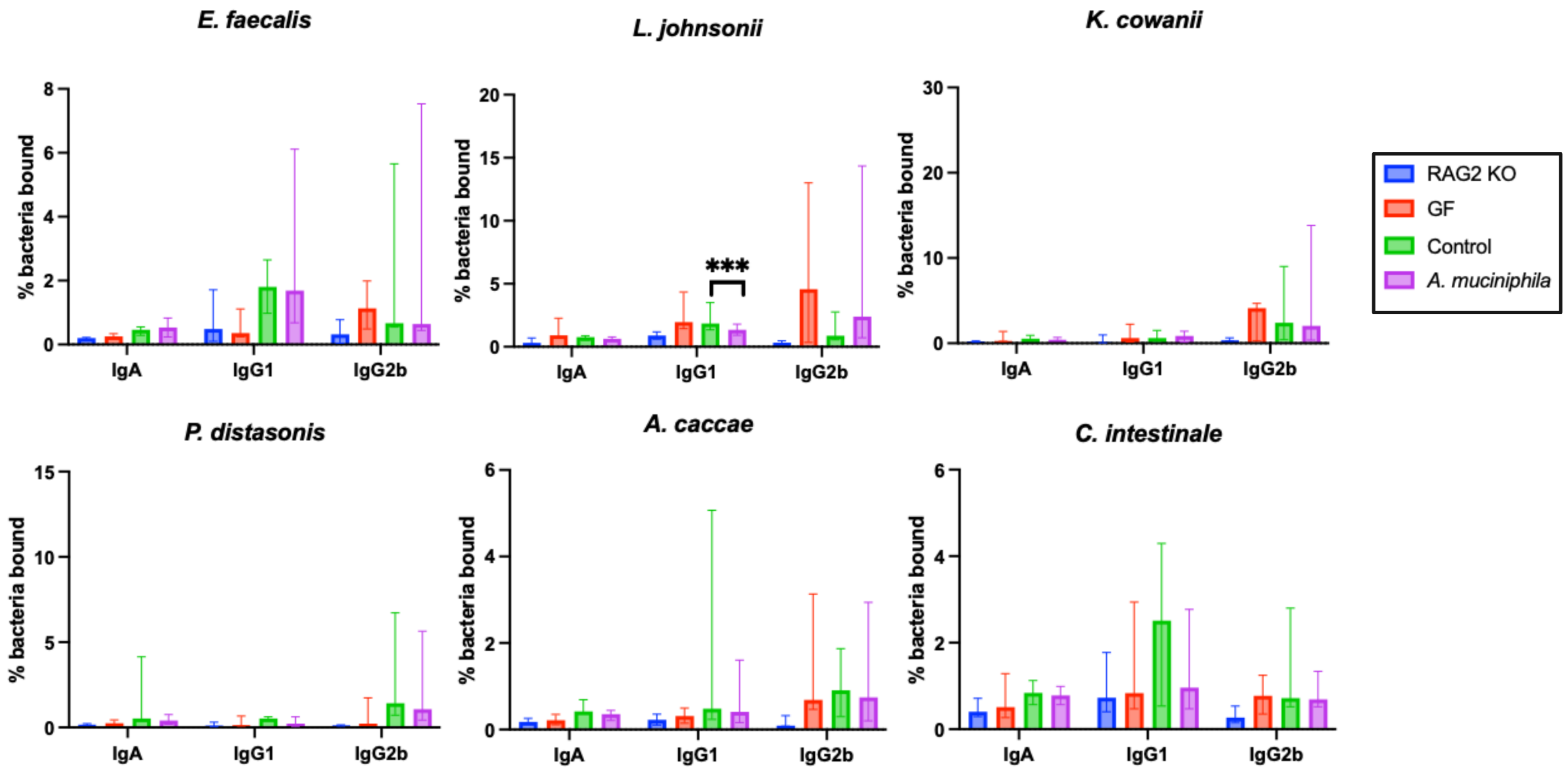

Figure S4. Antigen cross-reactivity does not fully explain *A. muciniphila* mediated enhancement of antibody responses to PedsCom staphylococcal species.

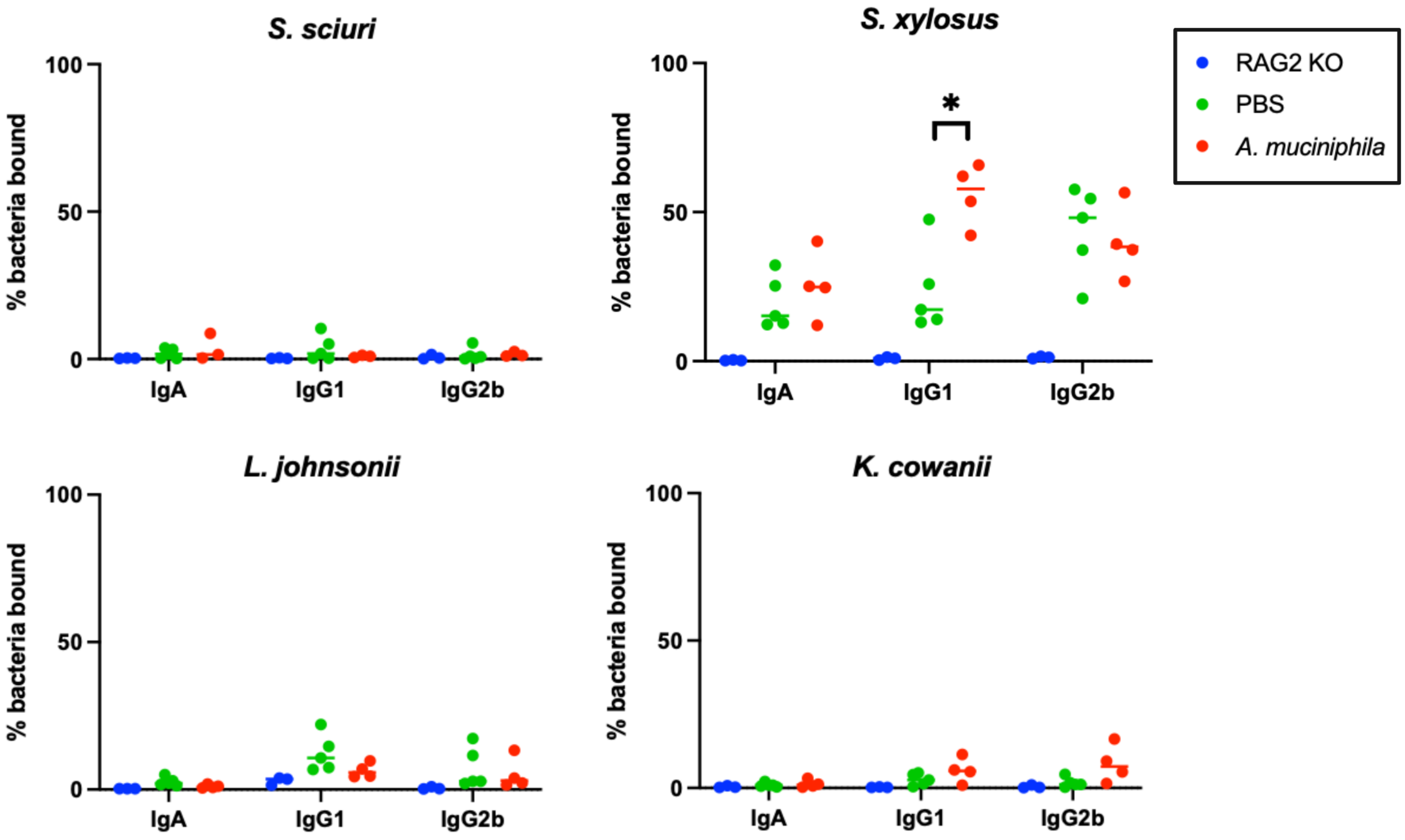

Figure S5. Vertically transferred *A. muciniphila* does not induce RORγ+ Tregs in PedsCom C57BL/6 mice.

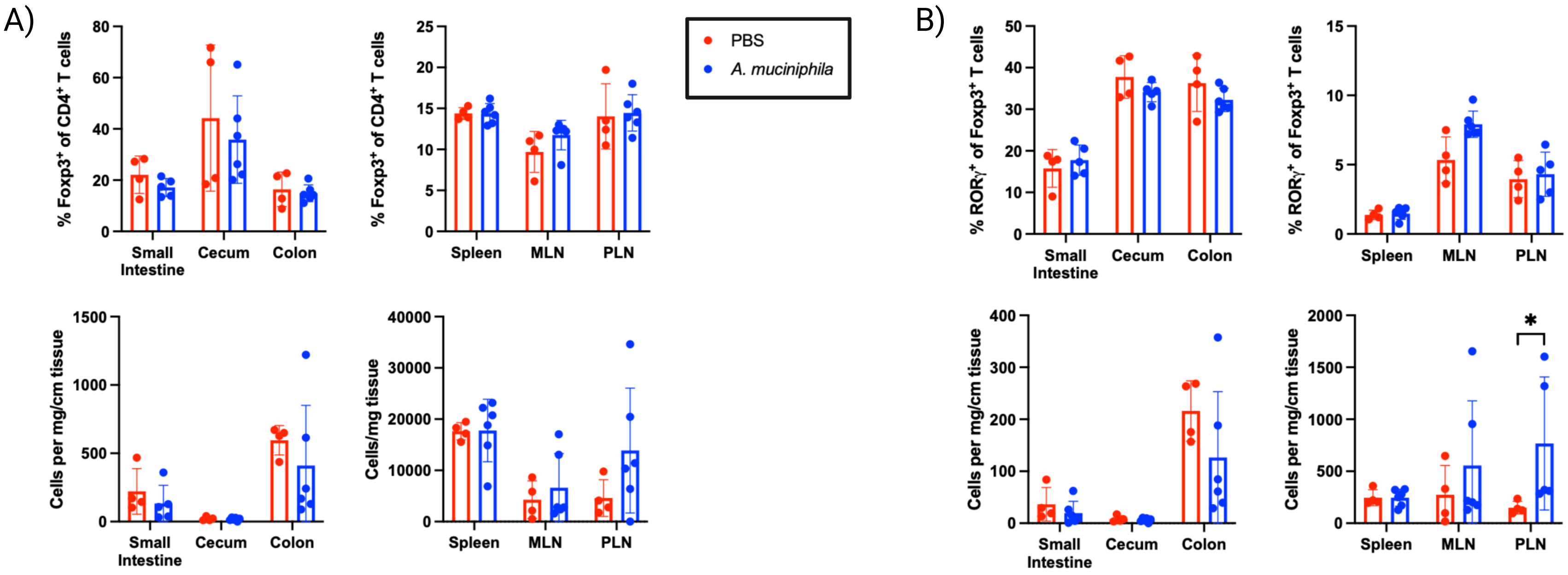

Figure S6. Vertically transferred *A. muciniphila* does not alter antibody responses to PedsCom microbes in C57BL/6 mice.

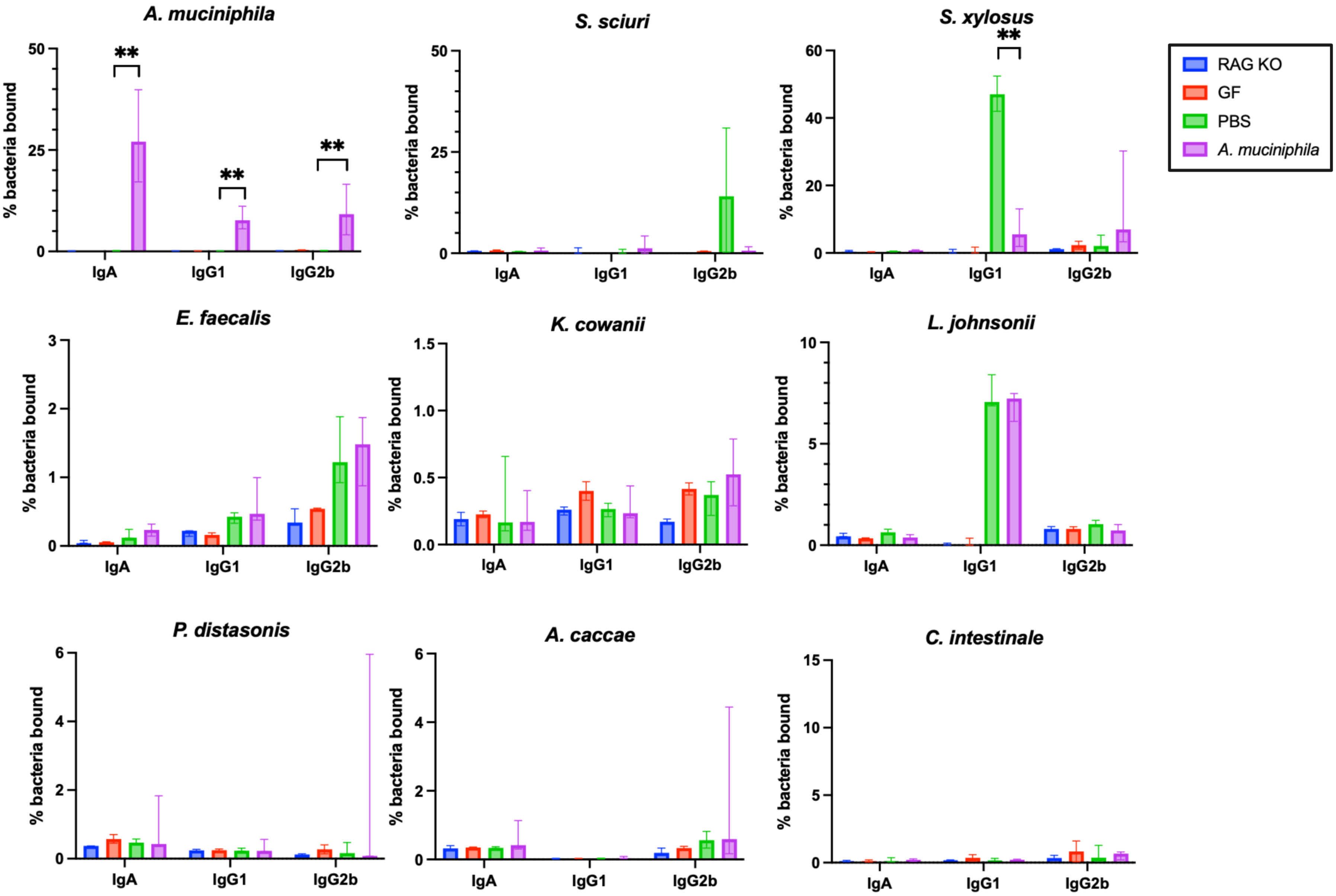
